# Evaluation of the novel culture-based FAST-T system allowing selection of optimal antibiotics for critically ill patients within 4 h (for other than bloodstream infectious)

**DOI:** 10.1101/519348

**Authors:** G Tetz, V Tetz

**Affiliations:** TGV-Biomed, New York, NY, USA; Human Microbiology Institute, New York, NY, USA

## Abstract

Rapid diagnostic tests are needed to improve patient care, particularly in immunocompromised hosts. Here, we describe the validation of a new phenotypic culture-based FAST-T method for rapid selection of antibiotics *in vitro* using specimens with mono-and polybacterial infections. FAST-T approach, which can be applied to any type of non-blood tissue, does not require isolation of pure bacterial cultures. FAST-T-selected antibiotics are those that can completely eliminate mixed bacterial infections in specimens. The method uses a novel FASM-T medium that allows more rapid bacterial growth of gram-positive and gram-negative monoisolates compared with that achieved with conventional laboratory media. The application of the FAST-T method in 122 bacterial species demonstrated overall sensitivity, specificity, positive predictive value, and negative predictive value of 99.6%, 98.1%, 98.5%, and 99.4%, respectively, already after 4 h. The overall category agreement with the outcome of standard testing was 98.9% with very major errors and major errors being detected in 1.2% and 0.6% of cases.

The use of FASM-T medium in 20 clinical polymicrobial samples allowed culturing a more diverse set of bacteria, including fastidious species, compared with that achieved with the standard laboratory diagnostic and enabled, already within 4 h, accurate selection of the antibiotics that completely eliminated all cultivatable bacteria from clinical samples. In conclusion, FAST-T system may be a valuable tool in improving phenotypic-based antibiotic selection, enabling targeted empirical therapy and accurate antibiotic replacement, which is especially important in high-risk patients.

## Introduction

Antibiotic therapy is typically started on an empirical basis, because the causative organism is not identified in an appreciable proportion of patients (1-3). However, empirical antibiotic therapy is inadequate in over 25% of cases, with 8–12% of patients receiving antibiotics that are ineffective (4). For these patients, antibiotic therapy must be adjusted following antimicrobial susceptibility testing (AST) by culture-based and/or molecular biology methods. However, even following the switch of the empirically chosen antimicrobial, in over 10% of cases, the newly selected antimicrobial remains ineffective (3). The resulting clinical failure is particularly dangerous for immunocompromised patients.

There are several types of diagnostic tests used currently for the selection of appropriate antibiotics, but neither of them is ideal with regard to its expedience and accuracy (5). Currently, traditional methods based on the isolation of bacterial culture, require at least 24 h to detect the growth of bacteria isolated from clinical specimens after sampling and another 6 to 24 h for detailed isolate characterization (i.e., biochemical identification) or susceptibility testing, during which time the treatment can only be empirical (6, 7). Therefore culture-based methods, owing to the slow turnaround time of results, are not suitable for tailored empirical therapy and are suboptimal in acute infection cases.

Another concern with culture-based tests is that media used in classical AST techniques, such as LB, Columbia and others, do not allow the growth of complex polymicrobial communities (8, 9). As a result, some clinically relevant organisms that are more difficult to culture do not get detected by standard AST of biological samples, leading to therapy failure (9). Many contemporary methods allow isolation of only the predominant species, whereas bacteria at the site of infection are rarely represented by a single species (10). The vast majority of the infections of the respiratory tract, urinary tract, skin, and soft-tissues had been previously characterized as being monomicrobial due to inaccurate results from culture-dependent isolation techniques. In contrast, the use of more advanced methods today indicates their polymicrobial nature (8, 11-14). Bacteria within polymicrobial communities are significantly more resistant to antimicrobials as they share collective antibiotic resistance (15). Furthermore, they are characterized by combined virulence contributing to the increased mortality rate among critically ill and compromised patients, such as patients with HIV or those that underwent solid organ or marrow transplantation (10, 16-18).

Another reason for the inappropriate selection of antibiotics by the standard AST methods is that they focus only on well-known pathogenic bacteria (8, 19). However, many bacteria that are considered nonpathogenic in non-immunocompromised patients may be pathogenic in people with suppressed immune response (20). This may happen because even in the case of successful elimination of one specific pathogen with a selected antibiotic, other bacteria that remain at the site of infection continue to grow and result in disease progression. Moreover, there is no benchmark accurate enough to indicate with certainty that bacteria are definitely pathogenic (21). For example, *Pandoraea apista, Aeromonas spp*., and *Kluyvera spp*. are now classified as pathogenic, although several decades ago, they were believed to be opportunistic and safe microorganisms (21-24). The media used in standard AST mostly promote the growth of the predominant bacteria and do not necessarily support proliferation of minor bacterial species from biological samples that may depend on each other for growth (25). That is why, it has been estimated that less than 10% of bacteria can be cultured in laboratory conditions and assayed by conventional AST (26-28).

In order to overcome these obstacles, in the last few decades, several types of novel diagnostic tests for rapid identification of bacteria and AST have been developed (29, 30). Such tests are based on direct pathogen identification using nucleic acid amplification techniques (5, 31). Although these methods are frequently utilized to determine the causative agents of bloodstream infections, some of them are used for the infections of other parts of the body (32-34).These methods do not depend on *in vitro* growth for bacterial detection and have become routine in many clinical microbiology laboratories. The introduction of PCR assays and sequencing technologies shortened the turnaround time to obtain the results and enabled characterization of pomyicrobial infections (35, 36). However, these assays also have a number of limitations. In particular, although the identification of bacteria helps to determine the therapeutic strategy, it does not really inform on the extent of resistance to antibiotics (6). Moreover, the methods based on 16S RNA sequencing are not very accurate as many bacterial species share similar or identical 16S rRNA sequences (37, 38).

Some molecular biology methods are based on the detection of antimicrobial resistance genes (39, 40). However, the reliance on the presence of resistance genes may be misleading with regard to accurate antibiotic selection as not all resistant genotypes result in resistant phenotypes (6). Furthermore, these methods can only detect known antibiotic resistance genes but may omit those, which have not yet been discovered (39, 41). The available resistance markers are not sufficiently comprehensive to provide clinically actionable results and may lead to the usage of either unnecessarily broad spectrum antibiotics or those without the sufficiently strong therapeutic effect.

Consequently, culture-based AST that predicts not only the resistance but also susceptibility to different drugs is still considered the gold standard for the identification of the most appropriate antibiotic for various infections (42).

Here, we evaluate a newly developed culture-based FAST-T system that can provide rapid phenotypic-based antibiotic selection for monobacterial and polybacterial clinical isolates. The FAST-T approach is based on a novel paradigm for the selection of effective antibiotics that considers not only monobacterial infection but also polybacterial cooperative interactions at the infection site.

## Materials and Methods

### Study samples and laboratory settings

Study specimens were obtained from the Human Microbiology Institute (New York, NY, USA), maintained at about 4 °C (wet ice), but not frozen, and sent to TGV-Biomed within 8 h after sampling.

Bronchoalveolar lavage (BAL) or sputum samples obtained from patients with chronic obstructive pulmonary disease (COPD), community-acquired pneumonia (CAP), ventilator-associated pneumonia (VAP), hospital-acquired pneumonia (HAP), cystic fibrosis (CF) were tested by using both state-of-the-art and FAST-T system methods.

### Antibiotics

AST was limited to antibiotics used for the treatment of respiratory infections, which were selected according to the American Thoracic Society/Infectious Disease Society of America guidelines (43-45). Antibiotics were added to FASM-T medium at respective maximal concentrations that could be achieved at the site of infection according to literature data: amikacin, 9 μg/mL (46); azithromycin 8 μg/mL (46, 47); aztreonam 22 μg/mL (48, 49); cefepime 24 μg/mL (46, 49, 50); gentamicin 5 μg/mL (56, 51); levofloxacin 12 μg/mL (46, 52); linezolid 25 μg/mL (53, 54); meropenem 10 μg/mL (55); piperacillin-tazobactam (25 for piperacillin and 3.5 for tazobactam) μg/mL (56); vancomycin 12 μg/mL (46, 57) (all from Sigma-Aldrich, St. Louis, MO). Probes were incubated at 37 °C, for 4, 8, and 24 h.

### Bacterial strains and growth conditions

Bacterial clinical isolates included the following gram-positive and gram-negative bacteria: *Staphylococcus aureus* (*n* = 22), *Streptococcus pneumoniae* (*n* = 4), *Streptococcus epidermidis* (*n* = 4), *Streptococcus pyogenes* (*n* = 2), *Enterococcus faecalis* (*n* = 5), *Bacillus cereues* (*n* = 4), *Paenibacillus VT400* (*n* = 1), *Bacillus respiratorii* VT-16-64 (*n* = 1) *Escherichia coli* (*n* = 15), *Acinetobacter baumannii* (*n* = 2), *Stenotrophomonas maltophilia (n = 4), Klebsiella pneumoniae* (*n* = 10), *Proteus vulgaris (n = 4), Proteus mirabilis* (*n* = 5), *Haemophilus influenzae* (n = 1), *Klebsiella oxytoca* (*n* = 1), *Rothia mucilaginosa (n = 1), Moraxella catharrhalis* (*n* = 1) *Pseudomonas aeruginosa* (*n* = 16), *Burkholderia cenocepacia* (*n* = 6), *Enterobacter cloacae complex* (*n* = 6), *Achromobacter xylosoxidans* (*n* = 2), *Serratia marcescens* (*n* = 3).

Bacteria were obtained from the Cystic Fibrosis Foundation Therapeutics Development Network Resource Center for Microbiology at the Seattle Children’s Hospital (Seattle, WA, USA) and from the Human Microbiology Institute (New York, NY, USA). Individual patterns of resistance to antibiotics are summarized in Table S1.

All bacterial strains were subcultured from frozen stocks onto Columbia broth (Oxoid Ltd., London, England) and incubated at 37 °C overnight. A standard bacterial inoculum of 5 × 10^5^ CFU/mL was used. All subsequent liquid subcultures were derived from colonies isolated from these plates and were further grown at different solid media: LB agar (Oxoid Ltd., London, England), Columbia agar (Oxoid Ltd., London, England), *Burkholderia cepacia*-selective agar (Hardy Diagnostics, Santa Maria, CA), chocolate agar (Oxoid UK), CHROMagar Staph aureus Medium (Becton, Dickinson and Company, Franklin Lakes, NJ), FAST-M medium (Human Microbiology Institute, New York, NY, USA). All media were supplemented with 5% lysed sheep erythrocytes (Becton Dickinson, Heidelberg, Germany).

### Biological specimen processing and bacterial isolation from them

BAL or sputum biospecimens (1 mL) from the patients with ventilator-associated, community-acquired pneumonia, cystic fibrosis (CF), or chronic obstructive pulmonary disease were mixed with 1 mL of sterile H_2_O until homogeneity with a plastic swab. After homogenization, 25 μL of the suspension was directly plated onto FASM-T medium in each well and incubated at 37 °C, for 4, 18, and 24 h. For control arm, probes were plated on LB agar and *Burkholderia cepacia*-selective agar (Hardy Diagnostics), chocolate agar (Oxoid), CHROMagar Staph aureus Medium (Becton, Dickinson and Company) and cultured according to laboratory recommendations at 37 °C for 24 h (8).

### Identification of bacteria in biological specimens

Bacterial identification to the species level from biological specimens was done by using subcultures on FAST-M or LB medium (Oxoid, UK). Isolates were examined for purity by light microscopy (Leica 2500DM, Leica Microsystems, Germany). To exclude the presence of mixed bacterial cultures, the isolates were assessed from at least 10 fields of view (59). After the subcultivation of every mixed bacterial culture up to monocultures, subsequent biochemical identification and matrix assisted laser desorption/ionization-time of flight (MALDI-TOF) mass spectrometry (Microflex LT, Bruker Daltonics, Germany) analysis were performed according to the manufacturer’s instructions after 24 h of growth.

The complete 16S rRNA gene sequencing of bacterial colonies was also used to identify the isolates. A PCR was performed with the general bacterial primers 27f (5′-AGAGTTTGATCCTGGCTCAG-3′) and 1492r (5′-GGTTACCTTGTTACGACTT-3′). The PCR mixture contained 0.2 μM of each primer and 40 ng of bacterial DNA in a total volume of 100 μL (HotStar Taq Master Mix Kit, Qiagen, Valencia, CA, USA) (60).

PCR was performed with a Mastercycler EP S gradient thermocycler (Eppendorf, Germany) using the following protocol: one cycle of 15 min at 95 °C, followed by 30 cycles, of (60 s at 95 °C, 45 s at 50 °C, and 90 s at 72 °C), and the final extension cycle of 10 min at 72 °C. The PCR products were verified by electrophoresis in 0.8% agarose gel, and the rest of the sample was purified and concentrated in 30 μL of demineralized water using a Qiaquick PCR purification kit (Qiagen) (61).

PCR amplicons were sequenced with a BigDye Terminator v1.1 cycle sequencing kit (Life Technology, Foster City, CA, USA). Identification to the species level was conducted by comparing the obtained sequences with those in BLAST database (http://blast.ncbi.nlm.nih.gov/Blast.cgi). Identifications to the genus and species-level were done on the basis of ≥97% and 99% identity of the 16S rRNA gene sequence to the reference, respectively (62).

### FAST-T system and interpretation of results

FAST-T system is a 12-well plate (TGV-Biomed, USA) filled with FASM-T medium that comprises pancreatic digests of casein, peptic digest of meat, heart pancreatic digest, yeast extract, starch, and water (Human Microbiology Institute, New York, NY, USA).

Clinical specimens were directly plated onto FASM-T medium in each well with a sterile swab, avoiding scratching or damaging the agar. Ten of the twelve wells contained FASM-T nutrient medium with antibiotics (one antibiotic per well) at concentrations that can be practically achieved at the site of infection. Two control wells contained antibiotic-free FASM-T medium. Plate reading was performed following sampling and incubation at 37 °C, for 4, 8, and 24 h.

The presence of microbial growth was identified with naked eye and confirmed with a stereoscopic microscope (Leica S6, Leica Microsystems, Germany). The signs of early bacterial growth, namely increased turbidity, hemolysis, appearance of film, and microcolonies, in the wells with antibiotics were compared with bacterial growth signs in control wells.

Microbial growth in any well meant that in the pathological material, there were microorganisms resistant to the particular antibiotic in the nutrient medium in this well. Therefore, such antibiotic was categorized as “ineffective”. Absence of bacterial growth in the well meant that the antibiotic present in the well killed all bacteria in the pathological material and was thus categorized as “effective”. In rare cases, there was no growth in control wells after 4 h, so those cultures were not analyzed further as is separately described in the Results below.

### Gold standard definition

Culture-based *in vitro* AST was selected as the standard of care method. The minimal inhibitory concentrations (MICs) for antimicrobials were determined by the broth microdilution method according to the Clinical and Laboratory Standards Institute (CLSI) guidelines (63, 64). The isolates were categorized as susceptible or resistant according to CLSI breakpoint guidelines with the only modification that “intermediate” isolates were treated as “resistant” (65). A standard bacterial inoculum of 5 × 10^5^ CFU/mL was used. Serial twofold dilutions of the antimicrobials were prepared in cation-adjusted Mueller-Hinton broth. MIC was defined as the lowest concentration of antibiotic that completely inhibited visible growth. Experiments were conducted in triplicate.

### Data analysis

The FAST-T system antibiotic selection results were compared to those obtained with culture-based gold standard microdilution method as described above. For the purpose of data analysis, the following definitions were used. A true-positive result occurred when both the FAST-T and standard-of-care methods indicated that the microorganism was resistant to the antibiotic (i.e., bacteria gave growth on the medium with antibiotic). A true-negative result occurred when both FAST-T and standard-of-care methods demonstrated sensitivity of the microorganism to the antibiotic (i.e., bacteria did not grow on the medium with antibiotic). A false-positive result occurred when the FAST-T system identified a strain as being sensitive to a certain antibiotic (organisms did not grow in FAST-T plates), but according to the standard-of-care method, the organism was resistant to that antibiotic, whereas a false-negative result was recorded when the FAST-T system suggested that the microorganism was resistant to the antibiotic, whereas the standard-of-care method indicated that it was sensitive. We calculated the values of accuracy, sensitivity, specificity, as well as positive predictive value (PPV) and negative predictive value (NPV) as previously described (66).

The total category agreement (CA) was determined and the results discrepant with those obtained by the state of care microdilution method were categorized (65). The very major errors were recorded in the cases when the FAST-T system indicated that an isolate was susceptible to the antibiotic, whereas according to the standard of care method, the isolate was resistant. The major errors were recorded in the cases when the FAST-T system suggested that an isolate was resistant to the antibiotic, whereas the standard of care method indicated that the isolate was susceptible. The rate of very major errors was calculated by dividing the total number of very major errors by the total number of strains determined as being resistant and multiplied by 100%. The rate of major errors was determined by dividing the total number of major errors by the number of strains determined as being susceptible and multiplied by 100%.

### Bacterial diversity analysis

The variety of bacteria that grew on the media used was characterized by the α-diversity indices such as non-parametric abundance-based coverage estimator (ACE) and Chao 1, indices which were managed and analyzed using R version 3.4.1 software (67).

Species richness (a count of different species), that gave growth on different media were represented on a dot plot, generated by package ‘ggplot2’ within R version 3.4.1 (67). All statistical analyses were conducted with a significance level of α = 0.05 (*P* < 0.05).

## Results

### Comparison of bacterial growth rate on FAST-T medium to that on other media

After 4 h of cultivation, visible growth was detected in 119 of 122 monomicrobial cultures (97.5%) in plates with FASM-T medium. Visible growth was not detected after 4 h only for *S. maltophilia* (*n* = 1), *S. pneumoniae* (*n* = 1), and *A. baumannii* (*n* = 1). However, these three strains became visible already after 8 h in culture (122/122; 100%).

Under the same conditions, visible growth on LB agar or Columbia agar after 4 h of culturing was detected for fewer microorganisms. In particular, culturing on LB agar revealed growth of only 69 out of 122 strains (56.5%), whereas 89 out of 122 strains (72.9%) grew on Columbia agar. Moreover, some *Streptococcus* spp did not reveal clear growth even after 8 h on these media and became visible only by 24 h (Fig. 1, Table S2).

**Figure 1:**
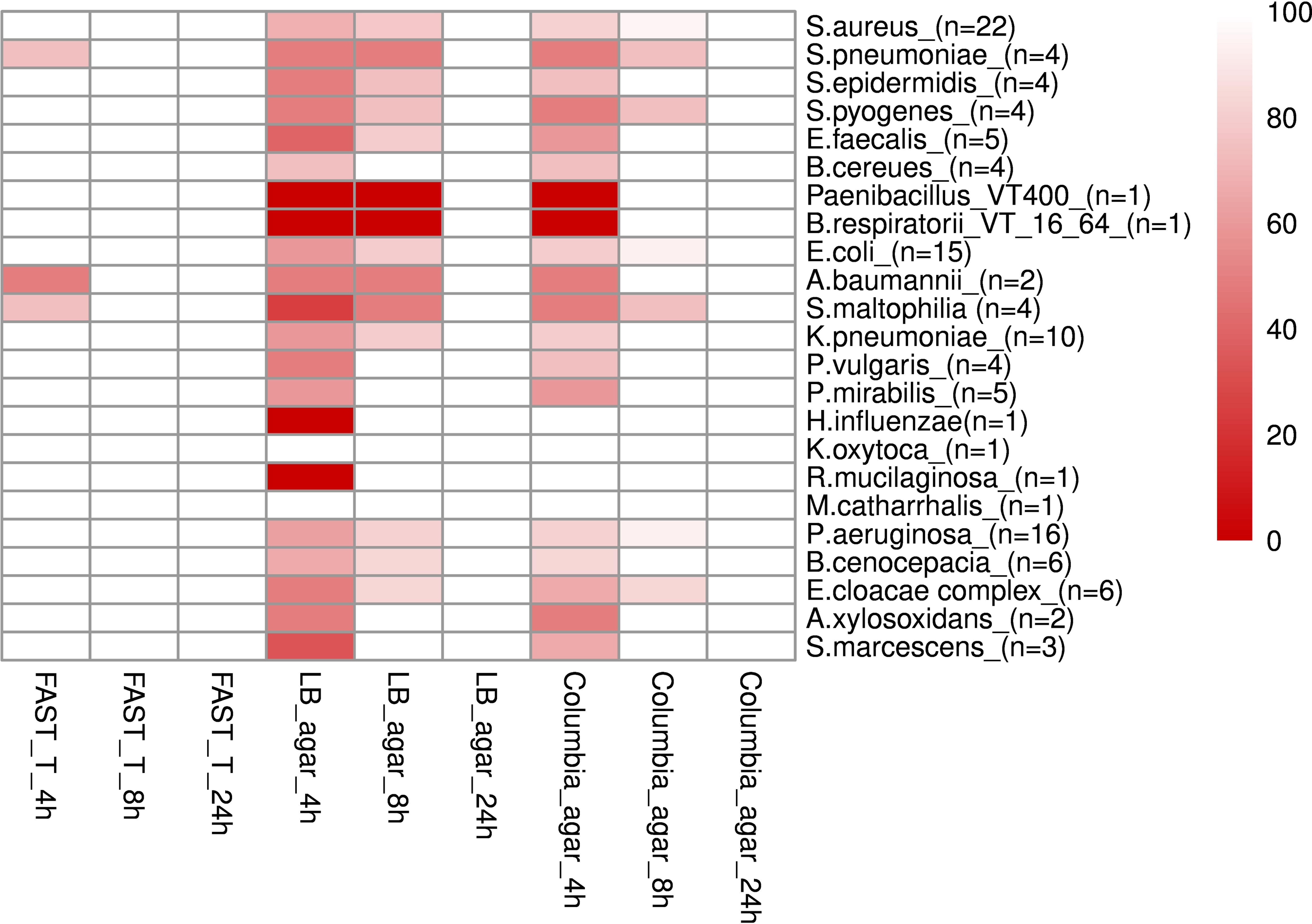
Comparison of bacterial growth rate on FASM-T, LB, and Columbia media. Growth rate is represented b y a heat map with each cell indicating the percentage of bacterial strains that gave growth at 4, 8, or 24 h. White color intensity represents the highest rate of growth, whereas red color indicates no growth for a particular time period.

We observed higher species richness after growth for 4 h on FASM-T medium as revealed by ACE and Chao 1 indices (28.07 and 33.5. respectively), compared with the values of these parameters after growth on LB agar (ACE □=□25.6 and Chao 1 =□21.5) or Columbia agar (ACE □=□25.5 and Chao 1 =□24; Fig. 2). Therefore, FASM-T was the only medium that allowed to detect visible growth of monomicrobial cultures already after 4 h with high accuracy.

**Figure 2:**
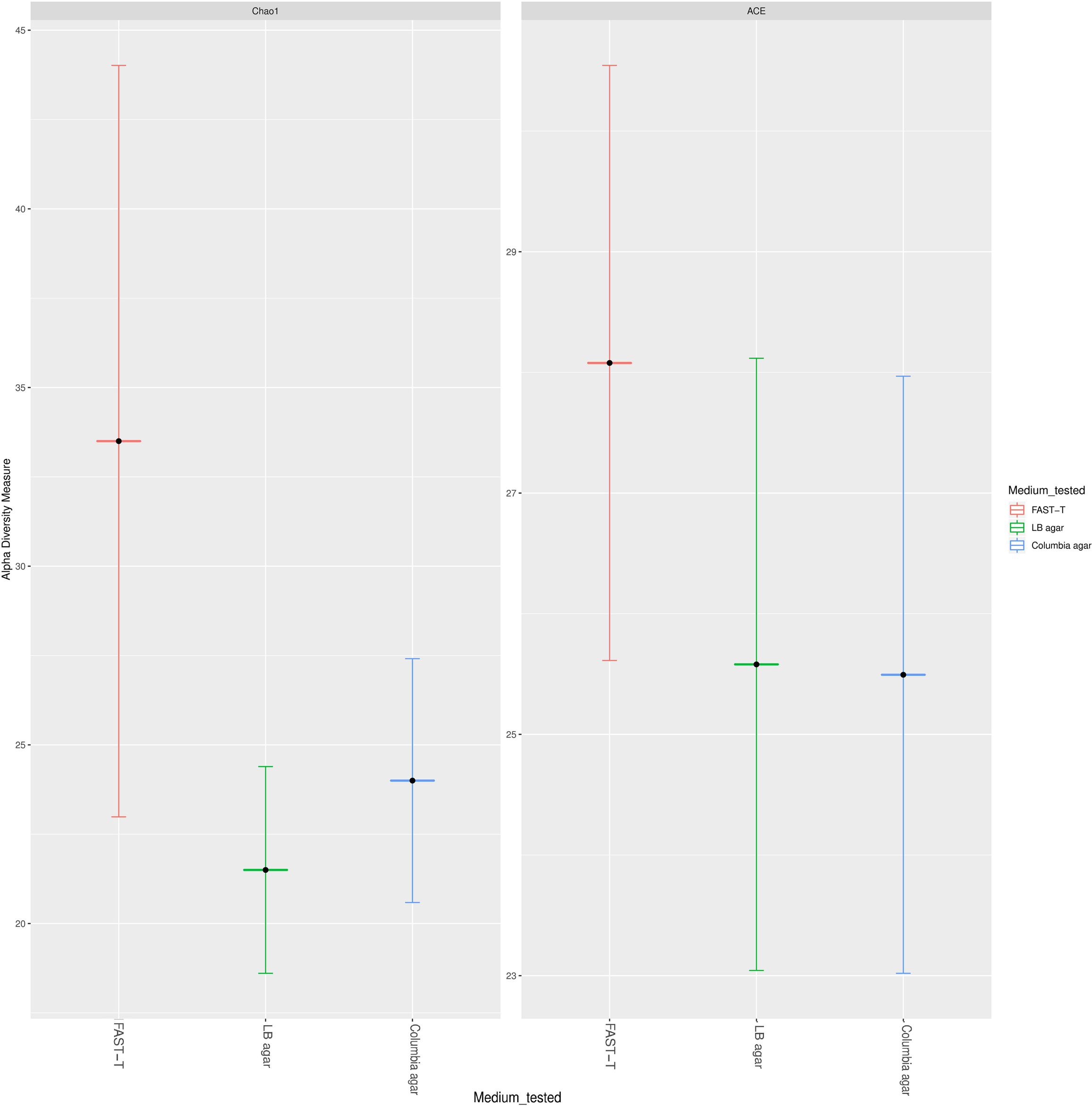
Increased diversity of bacterial species that gave growth on FASM-T medium, compared to that of species that grew on LB and Columbia agar, as revealed by the values of abundance-based coverage estimator (ACE), A) and Chao1 (B) indices.

### Estimation of the diversity of bacteria grown on FASM-T medium

Next, we applied FASM-T medium for the studies of polymicrobial infections. We used samples from patients with respiratory infections, which are usually associated with polymicrobial growth. Bacterial growth on FASM-T medium was seen within 4 h in 20 out of 20 clinical specimens (100%), whereas microbial growth was detected only in 8 out of 20 samples (40%) grown on LB agar that was used as standard medium. Only subsequent growth for 24 h revealed clear bacterial growth in all samples cultured in LB media.

Next, we analyzed polymicrobial growth in each sample by using FASM-T and LB media at 4 h and 24 h of culturing. A detailed description of bacteria that gave growth on different media is provided in Table 1. Already after 4 h, direct plating on FASM-T media allowed to cultivate mixed bacterial communities in 19 of 20 clinical specimens (95%). The increase of the culturing time to 24 h did not increase the diversity of cultured bacteria. Under the same conditions, no mixed communities (0/100; 0%) were observed during exposure to LB media after 4 h. Subsequent culturing for up to 24 h allowed to identify polymicrobial communities only in 5 out of 20 samples (25%).

**TABLE 1.** Comparison of diversity values of bacteria grown on FAST-T and standard media for 4 and 24 h.

| Number | Clinical specimen | Diagnosis | Bacteria identified with FASMT-T medium |  | Bacteria identified with standard media* |  |
| --- | --- | --- | --- | --- | --- | --- |
|  |  |  | 4 h | 24 h | 4 h | 24 h |
| 1 | BAL | COPD | *Burkholderia gladioli*<br>*Neisseria spp.*<br>*Haemophilus parainfluenzae*<br>*S. mitis*<br>*R.mucilaginoso*<br>*Streptomyces violaceruber* | *B. gladioli*<br>*Neisseria spp.*<br>*H.parainfluenzae*<br>*S. mitis*<br>*R.mucilaginoso*<br>*S. violaceruber* | - | *S. mitis*<br>*R. mucilaginoso* |
| 2 | BAL | COPD | *S.aureus*<br>*K.pneumonia*<br>*E. coli*<br>*E.faecalis*<br>*Proteus spp.* | *S.aureus*<br>*K.pneumonia*<br>*E. coli*<br>*E.faecalis*<br>*Proteus spp.* | *K.pneumonia* | *K.pneumonia* |
| 3 | BAL | CAP | *S.aureus*<br>*S.pneumonia* | *S.aureus*<br>*S.pneumonia* | - | *S.aureus* |
| 4 | BAL | CAP | *S.aureus* | *S.aureus* | *S.aureus* | *S.aureus* |
| 5 | BAL | CAP | *P.aeruginosa*<br>*H. influenza*<br>*M. catarrhalis* | *P.aeruginosa*<br>*H. influenza*<br>*M. catarrhalis* | *P.aeruginosa* | *P.aeruginosa* |
| 6 | BAL | VAP | *S.aureus*<br>*E.cloacae*<br>*P.aeruginosa* | *S.aureus*<br>*E.cloacae*<br>*P.aeruginosa* | - | *P.aeruginosa* |
| 7 | BAL | VAP | *S.aureus*<br>*K.pneumonia*<br>*E. coli*<br>*C.mucoviscidosis*<br>*S.aureus* | *S.aureus*<br>*K.pneumonia*<br>*E. coli*<br>*C.mucoviscidosis*<br>*S.aureus* | - | *S.aureus*<br>*E. coli* |
| 8 | Sputum | CF | *P.aeruginosa*<br>*S.aureus*<br>*R.mucilaginosa* | *P.aeruginosa*<br>*S.aureus*<br>*R.mucilaginosa* | *P.aeruginosa* | *P.aeruginosa*<br>*S.aureus* |
| 9 | Sputum | CF | *P.aeruginosa*<br>*S.aureus*<br>*S. maltophilia*<br>*B.cepacia* | *P.aeruginosa*<br>*S.aureus*<br>*S. maltophilia*<br>*B.cepacia* | - | *P.aeruginosa*<br>*B.cepacia* |
| 10 | Sputum | CF | *P.aeruginosa*<br>*Micrococcus luteus*<br>*S.aureus*<br>*S.oralis*<br>*S.milleri* | *P.aeruginosa*<br>*M.luteus*<br>*S.aureus*<br>*S.oralis*<br>*S.milleri* | - | *P.aeruginosa* |
| 11 | Sputum | CF | *P.aeruginosa*<br>*A.xylosoxidans*<br>*E.coli* | *P.aeruginosa*<br>*A.xylosoxidans*<br>*E.coli* | *P.aeruginosa* | *P.aeruginosa* |
| 12 | Sputum | CF | *S. maltophilia*<br>*E.coli*<br>*Bacillaceae spp*<br>*P.aeruginosa* | *S. maltophilia*<br>*E.coli*<br>*Bacillaceae spp*<br>*P.aeruginosa* | *P.aeruginosa* | *P.aeruginosa* |
| 13 | Sputum | CF | *P. aeruginosa*<br>*S. haemoliticus*<br>*A. baumannii*<br>*A. xylosoxidans*<br>*B. thailandensis* | *P. aeruginosa*<br>*S. haemoliticus*<br>*A. baumannii*<br>*A. xylosoxidans*<br>*B. thailandensis* | - | *P. aeruginosa* |
| 14 | Sputum | CF | *B. multivorans*<br>*Corynebacterium pseudodiphtherit*<br>*icum*<br>*Moraxella catharrhalis*<br>*Paenibacillus pabuli* | *B. multivorans*<br>*C.pseudodiphtheriticum*<br>*M.catharrhalis*<br>*P.pabuli* | - | *C. pseudodiphtheriticum* |
| 15 | Sputum | CF | *A. xylosoxidans*<br>*R.mucilaginosa*<br>*Elizabethkingia miricola*<br>*Microbacterium paraoxydans* | *A. xylosoxidans*<br>*R. mucilaginosa*<br>*E.miricola*<br>*M.paraoxydans* | - | *A. xylosoxidans* |
| 16 | Sputum | CF | *P. aeruginosa*<br>*S. aureus*<br>*E. coli*<br>*S.anginosus*<br>*B. sonorensis* | *P. aeruginosa*<br>*S. aureus*<br>*E. coli*<br>*S.anginosus*<br>*B. sonorensis* | *P. aeruginosa* | *P. aeruginosa* |
| 17 | Sputum | CF | *P. aeruginosa*<br>*S. aureus*<br>*E. coli*<br>*B. cereus* | *P. aeruginosa*<br>*S. aureus*<br>*E. coli*<br>*B. cereus* | *P. aeruginosa* | *P. aeruginosa*<br>*S. aureus* |
| 18 | BAL | HAP | *K. pneumonia*<br>*Rothia dentocariosa*<br>*A.ursingii*<br>*Bacillus pumilus* | *K. pneumonia*<br>*R. dentocariosa*<br>*A.ursingii*<br>*B. pumilus* | - | *K. pneumonia* |
| 19 | BAL | CAP | *H.influenzae*<br>*Actinobacillus suis*<br>*Eikenella corrodens*<br>*Actinomyces oris* | *H.influenzae*<br>*A. suis*<br>*E. corrodens*<br>*A.oris* | - | *H. influenzae* |
| 20 | BAL | COPD | *K. pneumoniae*<br>*H. influenza*<br>*Bacillus obstructivus* | *K. pneumoniae*<br>*H. influenza*<br>*Bacillus obstructivus* | - | *K. pneumoniae* |

Bacteria from all cystic fibrosis patient samples were plated on LB medium as well as cultivated on *Burkholderia cepacia* selective agar, chocolate agar, and CHROMagar Staph aureus medium.

Next, we determined the overall gain of identified microbial diversity that gave growth on FASM-T vs standard media, demonstrating that even after 4 h of culturing, FASM-T uncovered a more diverse set of microorganisms from the biological specimens compared with that revealed by the standard medium after 24 h of culturing (Fig. 3).

**Figure 3:**
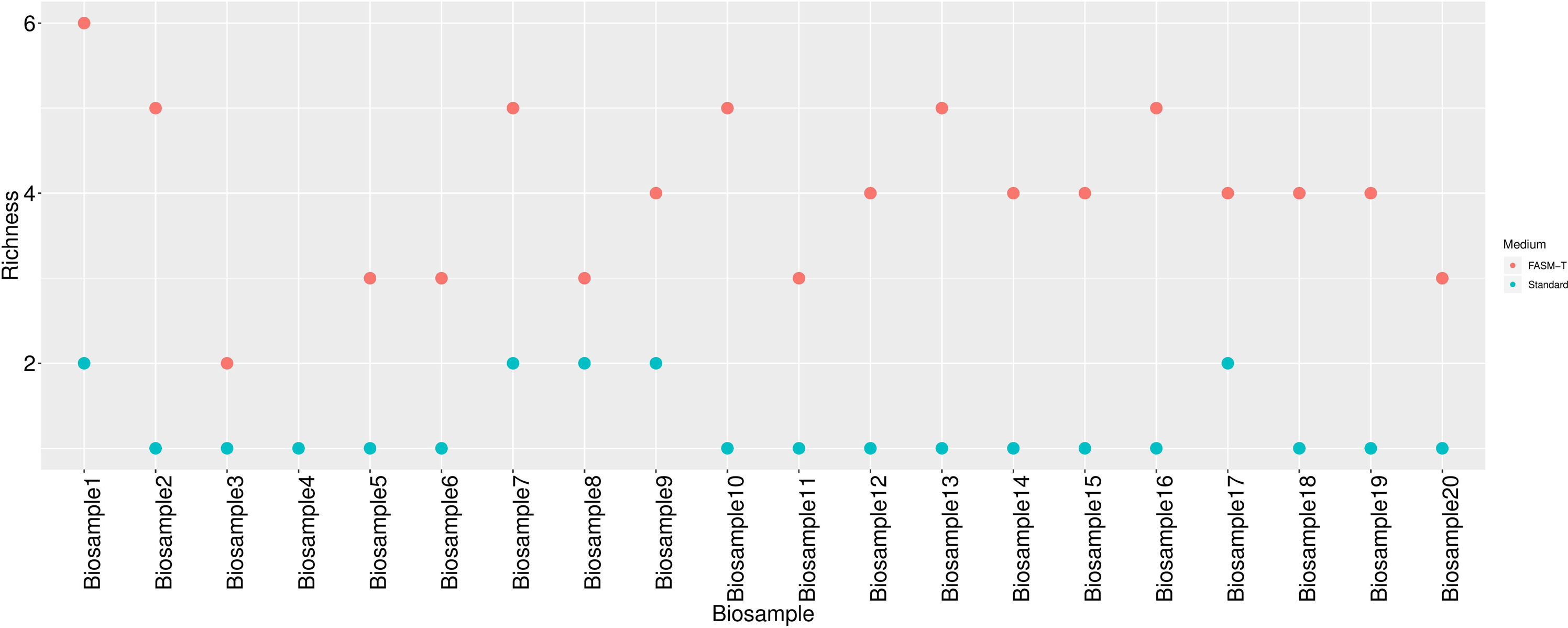
Comparison of the richness of bacteria that gave growth on FASM-T medium (after 4 h of culturing) compared with that of bacteria that gave growth (after 24 h) on standard media (LB medium, *Burkholderia cepacia*-selective agar, chocolate agar, CHROMagar Staph. aureus medium. Richness values for bio samples in two types of media are represented on a dot plot, displaying the distribution of numerical variables where each dot represents a value. The height of the column of dots represents the frequency for that value. The package ‘ggplot2’ within R (version 3.4.1) was used.

Notably, in the majority of specimens grown on FASM-T, we identified more than one well-known pathogen of respiratory tract infections, whereas the standard method allowed the identification of one and, only sometimes, two pathogens. The highest microbial diversity was observed in sputum of the patients with CF – a disease that is known to be characterized by a particularly complex, mixed lung microbiome (68). For example, from some tissue specimens, we isolated *P. aeruginosa, S. haemolyticus, A. baumannii, A. xylosoxidans,* and *B. thailandensis* with FASM-T medium. Each of these microorganisms has been shown to be associated with lung infections in CF, whereas the standard method identified only the dominant *P. aeruginosa* (69-71).

The ability of FASM-T to enable growth of fastidious species is illustrated by the isolation of *S. milleri* from CF patient #10. This bacterium was only recently associated with pulmonary complications in CF because previously, *S. milleri* could not be cultured on standard laboratory media (72, 73). Another notable example was the identification with FASM-T of *K. pneumonia, Rothia dentocariosa, Acinetobacter ursingii* and *Bacillus pumilus*, in sample #18 from the patient with hospital-acquired pneumonia (74). Each of these species had been shown previously to be associated with human infectinos. However, under the same conditions, the standard method identified only *K. pneumonia*. A similar trend was noted for all other tested specimens from cases of other respiratory diseases whereby FASM-T enabled identification of a wider range of known respiratory pathogens compared to that revealed by the standard method.

Notably, by using FASM-T medium, we identified two previously unknown bacterial species *Chryseobacterium mucoviscidosis* sp. nov. and *Bacillus obstructivus* sp. nov. (75, 76). These bacteria have not been identified by MALDI-TOF, as they had very low similarity to known species, and their further whole genome sequencing proved that they were previously unknown bacterial species highly enriched in genes encoding pathogenic and antibiotic resistance factors.

### Antibiotic selection in monomicrobial cultures using FAST-T method

The performance of the FAST-T system for direct selection of 10 different antibiotics prescribed for the treatment of respiratory infections caused by gram-negative or gram-negative bacteria was evaluated in a total of 122 monobacterial cultures after 4 h and 24 h of culturing. Cultures of 3 out of 122 strains (*S. pneumoniae* VT-SP-14, *S. maltophilia* VT-CFSM-4, and *A. baumannii* ATCC 17978) were excluded from the experiment due to the lack of growth in control wells after 4 h, leaving 1,990 runs in total for analysis (119 bacterial species tested against 10 antibiotics; 1 run = each antibiotic tested against one bacterial strain). For the tested 119 strains, correct antibiotic selection, as gauged by the results of conventional AST, was achieved after 4 h in 1,177 of 1,190 runs (98.9%). In rare cases, false-positive and false-negative results were obtained. The overall sensitivity, specificity, PPV, and NPV were 99.6%, 98.1%, 98.5%, and 99.4%, respectively (Tables 2 and S3).

**TABLE 2.** Overall identification performance of the FAST-T system after 4 and 24 h of culturing.

| Species | N true positive |  | N true negative |  | N false positive |  | N false negative |  | Sensitivity (%) |  | Specificity (%) |  | PPV (%) |  | NPV (%) |  |
| --- | --- | --- | --- | --- | --- | --- | --- | --- | --- | --- | --- | --- | --- | --- | --- | --- |
|  | 4 h | 24 h | 4 h | 24 h | 4 h | 24 h | 4 h | 24 h | 4 h | 24 h | 4 h | 24 h | 4 h | 24 h | 4 h | 24 h |
| *S.aureus* | 134 | 134 | 84 | 84 | 1 | 1 | 1 | 1 | 99,3 | 99,3 | 98,8 | 98,8 | 99,3 | 99,3 | 98,8 | 98,8 |
| *S.pneumoniae* | 12 | 16 | 18 | 24 | 0 | 0 | 0 | 0 | 100 | 100 | 100 | 100 | 100 | 100 | 100 | 100 |
| *S.epidermidis* | 16 | 17 | 23 | 23 | 1 | 0 | 0 | 0 | 100 | 100 | 95,8 | 100 | 94,1 | 100 | 100 | 100 |
| *S.pyogenes* | 10 | 10 | 30 | 30 | 0 | 0 | 0 | 0 | 100 | 100 | 100 | 100 | 100 | 100 | 100 | 100 |
| *E.faecalis* | 27 | 27 | 22 | 22 | 1 | 1 | 0 | 0 | 100 | 100 | 95,6 | 95,6 | 96,4 | 96,4 | 100 | 100 |
| *B.cereus* | 17 | 17 | 23 | 23 | 0 | 0 | 0 | 0 | 100 | 100 | 100 | 100 | 100 | 100 | 100 | 100 |
| *Paenibacillus* VT400 | 5 | 5 | 4 | 4 | 1 | 1 | 0 | 0 | 100 | 100 | 80 | 80 | 83,3 | 83,3 | 100 | 100 |
| *B.respiratorii* VT1664 | 3 | 3 | 7 | 7 | 0 | 0 | 0 | 0 | 100 | 100 | 100 | 100 | 100 | 100 | 100 | 100 |
| *E.coli* | 70 | 71 | 78 | 78 | 2 | 1 | 0 | 0 | 100 | 100 | 97,5 | 98,8 | 97,2 | 98,6 | 100 | 100 |
| *A.baumannii* | 8 | 16 | 1 | 4 | 0 | 0 | 0 | 0 | 100 | 100 | 100 | 100 | 100 | 100 | 100 | 100 |
| *S.maltophilia* | 19 | 28 | 10 | 11 | 1 | 1 | 0 | 0 | 100 | 100 | 90,1 | 91,6 | 95 | 96,6 | 100 | 100 |
| *K.pneumoniae* | 51 | 51 | 47 | 47 | 1 | 1 | 1 | 1 | 98,1 | 98,1 | 97,9 | 97,9 | 98,1 | 98,1 | 97,9 | 97,9 |
| *P.vulgaris* | 18 | 18 | 22 | 22 | 0 | 0 | 0 | 0 | 100 | 100 | 100 | 100 | 100 | 100 | 100 | 100 |
| *P.mirabilis* | 22 | 22 | 28 | 28 | 0 | 0 | 0 | 0 | 100 | 100 | 100 | 100 | 100 | 100 | 100 | 100 |
| *H.influenzae* | 2 | 2 | 8 | 8 | 0 | 0 | 0 | 0 | 100 | 100 | 100 | 100 | 100 | 100 | 100 | 100 |
| *K.oxytoca* | 6 | 6 | 4 | 4 | 0 | 0 | 0 | 0 | 100 | 100 | 100 | 100 | 100 | 100 | 100 | 100 |
| *R.mucilaginoso* | 4 | 4 | 6 | 6 | 0 | 0 | 0 | 0 | 100 | 100 | 100 | 100 | 100 | 100 | 100 | 100 |
| *M.catharrhalis* | 4 | 4 | 6 | 6 | 0 | 0 | 0 | 0 | 100 | 100 | 100 | 100 | 100 | 100 | 100 | 100 |
| *P.aeruginosa* | 122 | 122 | 35 | 35 | 2 | 2 | 1 | 1 | 99,2 | 99,2 | 94,6 | 94,6 | 98,4 | 98,4 | 97,2 | 97,2 |
| *B.cenocepacia* | 53 | 53 | 7 | 7 | 0 | 0 | 0 | 0 | 100 | 100 | 100 | 100 | 100 | 100 | 100 | 100 |
| *E. cloacae complex* | 31 | 31 | 29 | 29 | 0 | 0 | 0 | 0 | 100 | 100 | 100 | 100 | 100 | 100 | 100 | 100 |
| *A.xylosoxidans* | 16 | 16 | 4 | 4 | 0 | 0 | 0 | 0 | 100 | 100 | 100 | 100 | 100 | 100 | 100 | 100 |
| *S. marcescens* | 14 | 14 | 16 | 16 | 0 | 0 | 0 | 0 | 100 | 100 | 100 | 100 | 100 | 100 | 100 | 100 |
| Gram-positive bacteria | 403 | 408 | 259 | 265 | 6 | 5 | 2 | 2 | 99.9 | 99.9 | 96.8 | 97.2 | 97.4 | 97.9 | 99.6 | 99.6 |
| Gram-negative bacteria | 437 | 455 | 295 | 298 | 14 | 14 | 2 | 2 | 99.8 | 99.8 | 98.6 | 98,8 | 99.2 | 99.4 | 99.7 | 99.7 |

NPV, negative predictive value; PPV, positive predictive value.

After 24 h of culturing, we preformed 1,220 runs in total, as all bacteria at this time gave growth on FASM-T control antibiotic-free medium. By 18 h, all microbial strains gave growth in control wells. Correct antibiotic selection was achieved at 1,209 of 1,220 runs (99.1%). The values of analyzed parameters after 24 h of culturing were marginally higher than those achieved after 4 h (sensitivity, 99.6%; specificity, 98.5%; PPV, 98.8%; and NPV, 99.4%; Tables 2, S4). Notably, there was no difference in the overall ability to detect gram-positive and gram-negative bacteria. Next, we determined CA after 4 and 24 h of culturing for each antibiotic. After 4 h, the overall CA for all antibiotics used was 98.9% (1,177/1,190, 10 very major errors, 3 major errors). The rate of very major errors was 1.5% (10 very major errors/666 resistance outcomes), and the rate of major errors was 0.6% (3 major errors/512 susceptibility outcomes). The highest CA values were observed for levofloxacin, vancomycin (absolute agreement, 100%), followed by those for aztreonam, gentamicin, linezolid, piperacillin/tazobactam (for all: 118/119; 99.2%), amikacin, azithromycin, cefepime (for all: 117/119; 98.3%), and meropenem (116/119; 97.4%) (Table 3).

**TABLE 3.** Category agreement of antibiotic sensitivity results of the FAST-T method with those obtained by the standard method after 4 and 24 h of culturing.

| Antibiotic | Category agreement:<br>correct identifications/total No of<br>observations (%) |  | No of errors |  |  |  |
| --- | --- | --- | --- | --- | --- | --- |
|  |  |  | Very major<br>(false-positive) |  | Major<br>(false-negative) |  |
|  | 4h | 24h | 4h | 24h | 4h | 24h |
| Piperacillin-tazobactam | 118/119 (99,2) | 122/122 (100) | 1 | 0 | 0 | 0 |
| Meropenem | 116/119 (97,4) | 119/122 (97,5) | 2 | 2 | 1 | 1 |
| Levofloxacin | 119/119 (100) | 122/122 (100) | 0 | 0 | 0 | 0 |
| Aztreonam | 118/119 (99,2) | 121/122 (99,2) | 1 | 1 | 0 | 0 |
| Gentamicin | 118/119 (99,2) | 121/122 (99,2) | 0 | 0 | 1 | 1 |
| Amikacin | 117/119 (98,3) | 120/122 (98,3) | 2 | 2 | 0 | 0 |
| Azithromycin | 117/119 (98,3) | 121/122 (99,2) | 2 | 1 | 0 | 0 |
| Vancomycin | 119/119 (100) | 122/122 (100) | 0 | 0 | 0 | 0 |
| Cefepime | 117/119 (98,3) | 120/122 (98,3) | 2 | 2 | 0 | 0 |
| Linezolid | 118/119 (99,2) | 121/122 (99,2) | 0 | 0 | 1 | 1 |
| Overall performance | 1177/1190 (98,9) | 1209/1220 (99,1) |  |  |  |  |

CA values for the FAST-T system were even higher when the period of culturing was extended. The overall CA for all antibiotics after 24 h of cultivation was 99.1% (1,209/1,220) with 8 very major errors and 3 major errors. The rate of very major errors was 1.2% (8 very major errors/688 resistance results), and the rate of major errors was 0.6% (3 major errors/521 susceptibility results). CA values became nominally higher for piperacillin/tazobactam and azithromycin. Thus, after 24 h of culturing, the absolute CA was observed for piperacillin/tazobactam, levofloxacin, and vancomycin with no errors detected (100%). Slightly lower, but nonetheless very high CA values were noted for aztreonam, azithromycin, gentamicin, linezolid (for all: 121/122; 99.2%), amikacin, cefepime (for all: 120/122; 98.3%), and meropenem (119/122; 97.5%) (Table 3).

At both time points tested, the level of major errors was very low, indicating that antibiotic-sensitive isolates did not grow on FASM-T medium with antibiotics added at chosen concentrations. In general, there was no discernible pattern of the errors that could be explained by the nature of microorganisms or antibiotics tested: AST errors occurred in various species and with various drugs. CA values were not statistically different between the antibiotics used.

### Antibiotic selection in polymicrobial cultures using FAST-T method

We used 10 randomly selected samples from cases of respiratory infections tested before and analyzed particularities of antibiotic selection with the FAST-T (after growth for 4 h) and standard methods (Table 4). The majority of antibiotics identified by the standard method as “ineffective” (totally N50) were also identified as “ineffective” with FAST-T (total N65). At the same time, in several cases, there was a discrepancy between the conclusions of the FAST-T and standard methods about antibiotic efficacy. Some discrepancies were detected, in particular, for narrow-spectrum antibiotics, when growth of mixed gram-positive and gram-negative species was revealed by the FAST-T method, whereas only one species gave growth on the standard medium. However, such discrepancies were rare and that narrow-spectrum antibiotics were effective against mixtures gram-positive and gram-negative pathogens grown on FASM-T medium likely indicated complicated interspecies interactions (Table 6). For example, aztreonam was effective against mixed communities formed not only by the gram-negative *P. aeruginosa*, but also by *S. aureus*. Furthermore, vancomycin was found to be effective against mixed infections associated with both gram-positive and gram-negative bacteria.

**TABLE 4.**
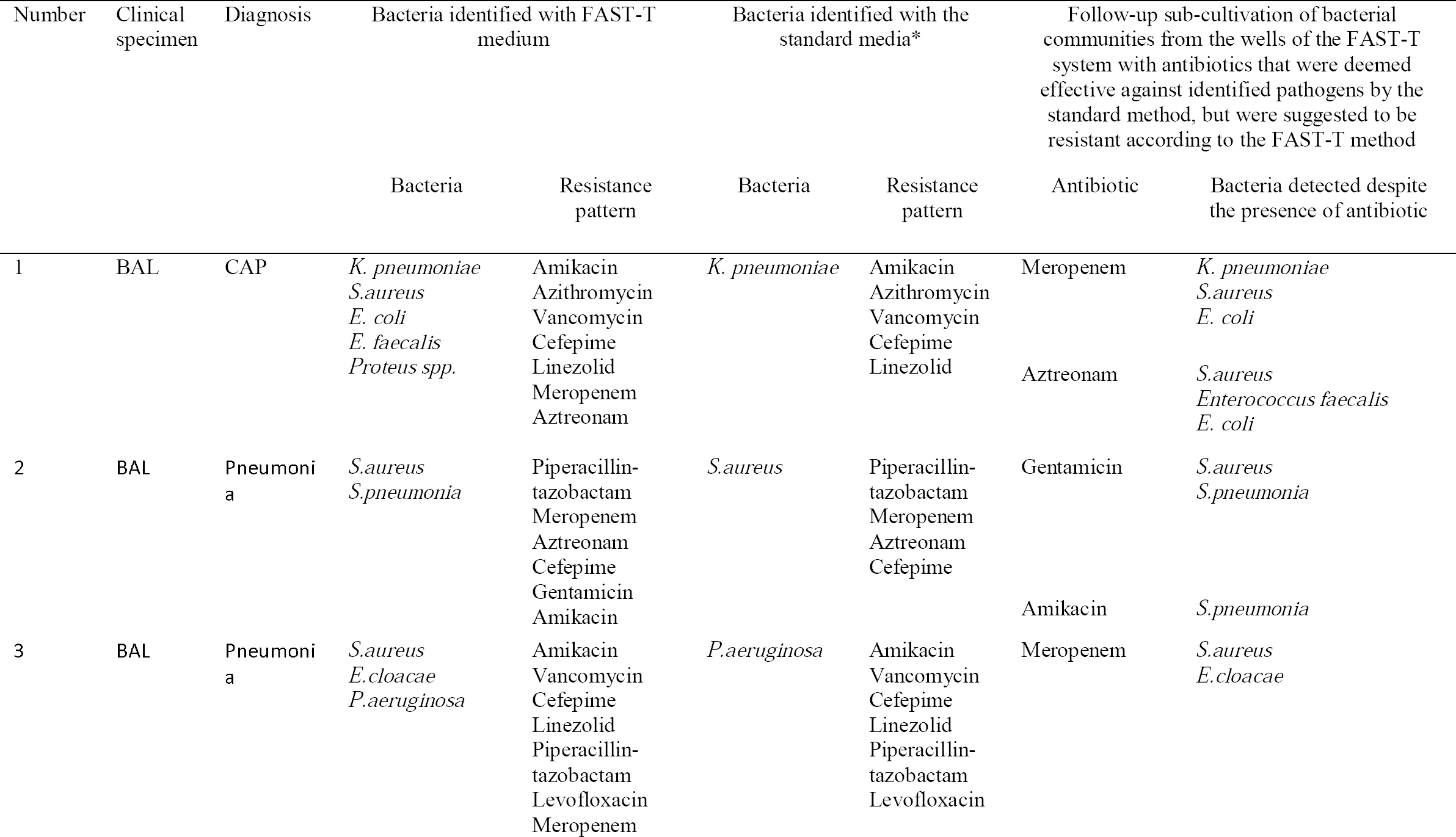
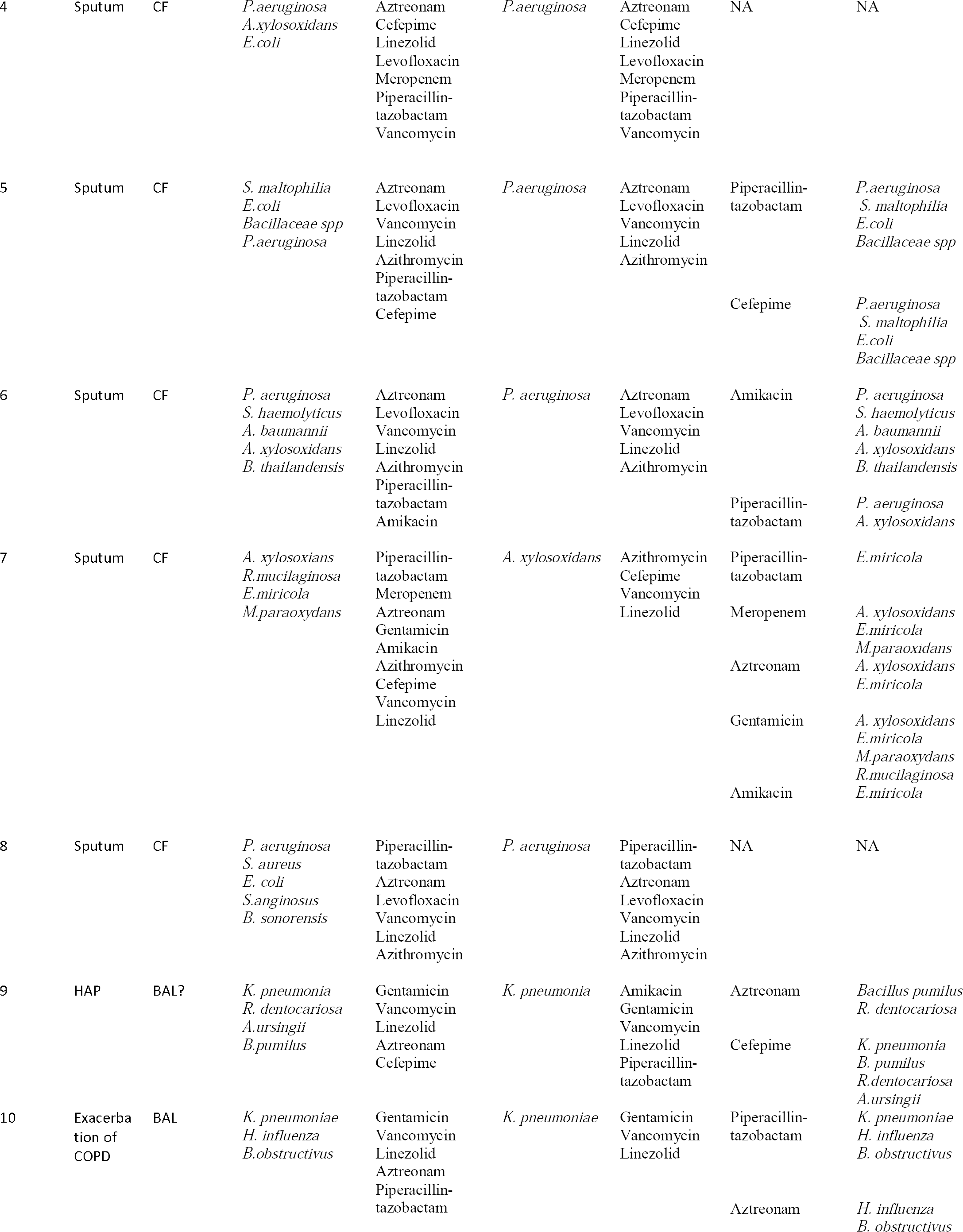
Identification of effective antibiotics selection from the biological samples with FAST-T (4 h of culturing) standard media (24 h of culturing).

We performed follow-up subculturing of bacterial colonies from the wells containing FASM-T with those antibiotics that were identified as being effective against the identified predominant pathogen by standard AST, but, as the presence of live colonies indicated, proved to be ineffective according to FAST-T. In 22% of such cases (11 discrepancies out of 50 cases), we could isolate the remaining identified predominant pathogen. Most likely this happened because complicated polymicrobial interactions at the site of infection that resulted in the higher resistance of bacteria, including the predominant pathogen, were not adequately simulated by the standard method compared to monobacterial culture growth. However, such interactions could be modeled well with the FAST-T system.

## Discussion

New diagnostic methods for the selection of antibiotics for tailored empirical therapy and for the change of antibiotic therapy that failed in immunocompromised patients are urgently needed. The availability of rapid and accurate methods of antibiotic selection for non-bloodstream infections, such as lung, urinary tract, skin, and soft tissue infections, would have a great impact on the disease outcome, length of hospital stay, possible complications, and spread of antibiotic resistance (77).

Our present experiments evaluated for the first time the performance of the novel FAST-T system providing culture-based antibiotic selection within a short, 4 h period. The FAST-T method can be used for initial tailored empirical therapy and for the selection of an appropriate antibiotic to those patients who have not responded to the previous therapy and require change of antimicrobial treatment strategy. Our original hypothesis that the use of the FAST-T system would provide a rapid and accurate selection of antibiotics for both monobacterial or polybacterial infections sampled directly from clinical specimens, was found to be confirmed experimentally. FAST-T system allowed faster growth of monobacterial cultures compared with that achieved by the standard method used in clinical diagnostics and demonstrated an increased richness with higher ACE and Chao 1 indices for bacteria that gave growth within 4 h on FASM-T medium than those recorded for bacteria grown on LBA or Columbia agar (Fig. 2).

Our study also revealed that FAST-T system provided accurate antibiotic selection compared to that achieved by the standard of care, with results being available already after 4 h of culturing. If compared with the standard method that involves the isolation of pure bacterial cultures prior to AST, the FAST-T system was faster by more than 32 h. The antibiotic selection was highly accurate already within 4 h (sensitivity, 99.6%; specificity, 98.1%; PPV, 98.5%; NPV, 99.4%, and CA, 98.9%); with no discernable bias in the error pattern toward gram-positive or gram-negative bacteria or toward a particular antibiotic class. Furthermore, the increase of culturing time from 4 h to 24 h did not significantly change the accuracy of the FAST-T method performance, indicating that accurate results can be obtained after just several hours of culturing.

In the current study, FASM-T medium supported growth of a significantly more diverse set of bacteria from biological samples even after 4 h of culturing than the standard medium, broadly used in microbiological laboratories, after 24 h of culturing. Thus, FAST-T system allowed identification of polymicrobial cultures in 19 out of 20 samples after 4 h of culturing. This was in agreement with previous reports that emphasized the polymicrobial nature of the majority of respiratory tract infections (13). Under the same conditions, the latter approach detected polymicrobial cultures in only 5 out of 20 samples. In this study, the standard method failed to detect not only some isolated commensal bacteria, but also well-known pathogens, such as *B. cepacia, S.maltophila, K. pneumonia*, and others.

Notably, many bacteria identified within the polybacterial samples are known as causative agents of respiratory infections. For example, we detected *S. aureus, R. dentocariosa, H. influenzae, E. corrodens, E. miricola, A. baumannii, A. xylosoxidans*, and other well-known respiratory tract pathogens in samples from CF and chronic obstructive pulmonary disease cases. These microorganisms have been shown to cause recurrent respiratory infections but are completely overlooked by standard of care diagnostics. In the current study, the FAST-T system enabled growth of different complex mixtures of bacteria without a limit to the number of species that could be isolated. This finding is of primary importance due to two considerations. First, the presence of polymicrobial infections is one of the reasons for the inappropriate antibiotic selection and mortality of high-risk patients. The elimination of the dominant strains leads to the re-growth of other bacteria at the site of the infection unaffected by the initial antibiotic treatment, which may lead to disease progression and complications, e.g., due to the lack of sufficiently strong immune response. Moreover, as it has been shown before, some bacteria that are considered nonpathogenic in the lungs of non-immunosuppressed subjects may be pathogenic in immunosuppressed hosts (78, 79). This is why, for some bloodstream infections, sterilization is widely used, particularly in the patients with impaired immune response. Therefore, the fact that FAST-T system reliably detects both major and minor bacterial pathogens within the site of infection makes it promising for the use in immunocompromised patients. Second, our assessment of antibiotic sensitivity within polymicrobial communities showed that in some cases, antibiotics selected with the standard method did not eliminate even the dominant bacteria in mixed microbial communities. Such false-positive data might result in the selection of ineffective antibiotics and therapy failure (80). The reason for false-positive results generated by the gold standard method may be because this method enables growth of monobacterial cultures but does not take into consideration higher tolerance to antibiotics of bacteria within mixed communities (15). However, “real life” infections are predominantly polymicrobial by nature (10). The resulting survival of the lead pathogen leads to its re-growth and therapy failure. According to our *in vitro* data, FAST-T system is less prone to such limitations.

Furthermore, the use of the FAST-T system for the treatment of polybacterial infections allowed selection of not only broad spectrum but also narrow spectrum antibiotics, confirming the presence of complexed interspecies interactions in mixed bacterial communities.

Notably, by using FAST-T, we have isolated previously unknown bacterial species *Chryseobacterium mucoviscidosis* sp. nov. and *Bacillus obstructivus* sp. nov. that possessed several typical virulence factors, such as hemolysins, alpha-amylases, and others, which are found in other respiratory pathogens (75, 76, 81). Moreover, in these bacteria, we identified several antibiotic resistance genes that once found in endospore-forming *Bacillus* spp. are of concern because of the possible spread of antibiotic resistance among sporobiota members (82). The identification of previously unknown bacteria as well as some fastidious species, such as *S. milleri*, indicates that the developed FASM-T medium provides unique growth factors and cultivation environment that are not achievable with the standard media (19, 83).

The use of the FAST-T system enables to quickly select antibiotics that are effective for the treatment of each particular condition (i.e., those that kill *all* microorganisms present in the clinical specimen) without the need to identify precisely the causative agents of the infection or to determine MICs. Importantly, although the FAST-T method does not allow immediate identification of bacteria on the species level, bacterial cultures that gave growth on FASM-T medium can be isolated and subsequently identified by using culture-based techniques and any other standard methods for the study of bacteria.

FAST-T system has all benefits of phenotypic culture-based methods, but is 30–54 h faster than standard culture-based diagnostics that requires time-consuming isolation of pure bacterial cultures. Even compared with direct AST, the FAST-T method allows antibiotic selection on a much shorter timescale (84). Like direct AST, FAST-T system enables direct sampling of biological specimens without the need for culturing or time-consuming sample processing. Furthermore, the FAST-T method can indicate the suitable antibiotic in only 4 h, whereas direct AST requires 18–36 h (and that is why direct AST is not applicable for tailoring empirical antibiotic therapy) (84).

Despite being as fast as some of the molecular biology methods, FAST-T system lacks main disadvantages of molecular methods based on next-generation sequencing and 16S RNA sequencing, such as overestimation of antimicrobial resistance and inability to inform on antimicrobial susceptibility (37, 38).

In this study, we used FAST-T system with a set of antibiotics used for the treatment of respiratory infections. However, it can easily be adjusted for the diagnosis of the infections of other parts of the body, such as urinary tract, skin, or soft tissues, by including the antibiotics used for the therapy of these infections in FASM-T medium. Moreover, it can be used for the cultivation of bacteria with different oxygen requirements. The presence of bacterial growth was analyzed with naked eye and stereoscopic microscopy, but a more sophisticated device for visual monitoring can be used to increase the analysis accuracy. We believe that FAST-T system may become a valuable tool in improving antibiotic selection in diagnostics, with as little as 4 h turnaround time. The use of FAST-T system in therapeutic management will allow faster development of adequate therapies, including targeted empirical therapy and accurate antibiotic replacement, especially in immunocompromised, high-risk patients. Future studies will be necessary to investigate clinical efficacy of the FAST-T system and the clinical impact of its use alone and as an auxiliary method for standard diagnostics.

## Acknowledgments

This research received no specific grant from any funding agency in the public, commercial, or not-for-profit sectors. VT and GT have applied for a patent for the FASM-T medium. We thank D. Shalaikin for bioinformatics analysis.

---

**Supplementary Table S1.** Individual patterns of resistance to antibiotics.

**Supplementary Table S2**. Comparison of growth speeds of monoisolates on different media.

**Supplementary Table S3**. Results of antimicrobial susceptibility testing by using the FAST-T system (4 h culturing) and standard method (24 h culturing) for 119 gram-positive and gram-negative pathogens.

**Supplementary Table S4**. Results of antimicrobial susceptibility testing by using the FAST-T system and standard method for 122 gram-positive and gram-negative pathogens after 24 h of culturing.

