## Supplementary table 2 for "Evaluation of the novel culture-based FAST-T system allowing selection of optimal antibiotics for critically ill patients within 4 h (for other than bloodstream infectious)"

**Supplementary** **Table S2**. Comparison of growth speeds of monoisolates on different media.

| Bacteria (total number of strains tested) | Number of strains that gave growth at the media | | | | | | | | |
| --- | --- | --- | --- | --- | --- | --- | --- | --- | --- |
|  | FAST-T | | | LB agar | | | Columbia agar | | |
|  | 4h | 8h | 24h | 4h | 8h | 24h | 4h | 8h | 24h |
| *S.aureus* (n=22) | 22 | 22 | 22 | 15 | 17 | 22 | 18 | 21 | 22 |
| *S.pneumoniae* (n=4) | 3 | 4 | 4 | 2 | 2 | 4 | 2 | 3 | 4 |
| *S.epidermidis* (n=4) | 4 | 4 | 4 | 2 | 3 | 4 | 3 | 4 | 4 |
| *S.pyogenes* (n=4) | 4 | 4 | 4 | 2 | 3 | 4 | 2 | 3 | 4 |
| *E.faecalis (n=5)* | 5 | 5 | 5 | 2 | 4 | 5 | 3 | 5 | 5 |
| *B.cereues* (n=4) | 4 | 4 | 4 | 3 | 4 | 4 | 3 | 4 | 4 |
| *Paenibacillus* VT400 (n=1) | 1 | 1 | 1 | 0 | 0 | 1 | 0 | 1 | 1 |
| *B.respiratorii* VT-16-64 (1) | 1 | 1 | 1 | 0 | 0 | 1 | 0 | 1 | 1 |
| *E.coli* (n = 15) | 15 | 15 | 15 | 9 | 12 | 15 | 12 | 14 | 15 |
| *A.baumannii*  (n = 2) | 1 | 2 | 2 | 1 | 1 | 2 | 1 | 2 | 2 |
| *S.maltophilia (n=4)* | 3 | 4 | 4 | 1 | 2 | 4 | 2 | 3 | 4 |
| *K.pneumoniae* (n = 10) | 10 | 10 | 10 | 6 | 8 | 10 | 8 | 10 | 10 |
| *P.vulgaris (n=4)* | 4 | 4 | 4 | 2 | 3 | 4 | 3 | 4 | 4 |
| *P.mirabilis* (n = 5) | 5 | 5 | 5 | 3 | 5 | 5 | 3 | 5 | 5 |
| *H.influenzae* (n=1) | 1 | 1 | 1 | 0 | 1 | 1 | 1 | 1 | 1 |
| *K.oxytoca* (n = 1) | 1 | 1 | 1 | 1 | 1 | 1 | 1 | 1 | 1 |
| *R.mucilaginosa* (n=1) | 1 | 1 | 1 | 0 | 1 | 1 | 1 | 1 | 1 |
| *M.catharrhalis* (n=1) | 1 | 1 | 1 | 1 | 1 | 1 | 1 | 1 | 1 |
| *P.aeruginosa* (n = 16) | 16 | 16 | 16 | 10 | 13 | 16 | 13 | 15 | 16 |
| *B.cenocepacia* (n=6) | 6 | 6 | 6 | 4 | 5 | 6 | 5 | 6 | 6 |
| *E. cloacae* complex (n = 6) | 6 | 6 | 6 | 3 | 5 | 6 | 4 | 5 | 6 |
| *A.xylosoxidans* (n=2) | 2 | 2 | 2 | 1 | 2 | 2 | 1 | 2 | 2 |
| *S. marcescens* (n = 3) | 3 | 3 | 3 | 1 | 3 | 3 | 2 | 3 | 3 |
